# HoloCell: A Generative Foundation Model for Holistic Cellular Modeling

**DOI:** 10.64898/2026.06.07.730684

**Authors:** Qun Jiang, Zhen Li, Bowen Hu, Yunlong Bie, Keyi Li, Qiuyi Li, Peiran Jin, Yuan He, Pan Deng, Zhenyi Wang, Xiaoyang Chen, Tao Qin, Haiguang Liu, Rui Jiang, Qijin Yin

## Abstract

Single-cell multi-omics technologies have recently advanced to enable the profiling of epigenomic, transcriptomic, and proteomic layers within individual cells, offering new opportunities to characterize cellular states as integrated biological systems. However, developing a unified framework that can seamlessly integrate diverse omics modalities and remain robust to heterogeneous modality missingness remains challenging. Existing methods are often designed for specific modalities or modality pairs, relying on dataset-specific training or paired measurements. Here we present HoloCell, to our knowledge the first generative foundation model for joint representation learning and generative modeling across all three major single-cell omics modalities, i.e., epigenomics, transcriptomics, and proteomics. HoloCell contains over 860 million parameters and is pretrained on the Human-Multi-Omics-Corpus, which comprises approximately 468 million single-cell profiles across these three omics layers, corresponding to over 425 billion tokens. HoloCell introduces a a simple yet biologically motivated hierarchical tokenization strategy that encodes cis-regulatory elements, genes, and proteins as structured tokens within a shared modeling framework. We evaluated HoloCell across single-omics representation learning, paired multi-omics integration, unpaired multi-omics alignment, and cross-modal generation via iterative diffusion and remasking, demonstrating its superior performance and flexibility across diverse omics tasks. From a representation perspective, HoloCell provides a unified digital mapping of cellular states across multiple omics layers, capturing cell heterogeneity as an integrated system. From a generation perspective, its iterative diffusion and remasking framework permits flexible generation orders beyond fixed left-to-right causality, enabling in silico simulation of multi-omics information flow. Together, these capabilities position HoloCell as a versatile foundation model toward the emerging concept of a virtual cell, offering both systematic characterization and generative simulation of cellular systems within a unified framework.

## 1 Introduction

Recent advances in single-cell omics technologies have fundamentally transformed our ability to characterize cellular heterogeneity at unprecedented resolution. High-throughput platforms for single-cell transcriptomics, epigenomics, and proteomics have generated datasets at rapidly increasing scale, enabling systematic interrogation of gene regulation, chromatin accessibility, and protein expression across diverse biological systems [1, 2, 3]. This accumulation of single-cell measurements provides an unprecedented opportunity to learn general principles of cellular organization.

Early computational methods for single-cell analysis were developed largely within individual modalities. In transcriptomics, probabilistic and deep generative models such as scVI [4] established powerful frameworks for representation learning and differential expression. For epigenomics, deep learning approaches like scBasset [5] and CASTLE [6] demonstrated effectiveness in modeling chromatin accessibility and regulatory sequence features. In the domain of protein abundance and cytometry data, workflows such as cytoVI [7], CyCombine [8], and FlowSOM [9] provided robust solutions for normalization, batch correction, and clustering. While these methods have substantially advanced modality-specific analysis and remain important baselines, the rapid development of multi-omics profiling technologies has made single-modality modeling increasingly insufficient for comprehensive cell representation. To integrate information across different omics layers, various multi-modal and multi-omics methods have been proposed, including Seurat v4 [10], totalVI [11], and scGLUE [12], which have proven highly effective for paired integration, batch correction, and joint embedding construction. In parallel, generative approaches for denoising, imputation, or cross-modal prediction have been developed, ranging from probabilistic models like scVI and totalVI to recent diffusion-based models for single-cell multi-omics generation [13]. These developments have broadened single-cell modeling toward integrative and generative frameworks, further motivating generalizable models capable of representing cellular states across diverse biological contexts and adapting to multiple downstream tasks.

Driven by the rapid progress of large-scale pretraining and the growing availability of large and diverse single-cell datasets, foundation models have recently emerged as a promising direction for single-cell biology. In transcriptomics, models such as scGPT [14], scFoundation [15], and Geneformer [16] demonstrate that pretraining on large-scale gene expression datasets can yield transferable representations for diverse downstream tasks. These models typically adapt language-modeling ideas to single-cell expression profiles by treating genes or gene-associated signals as tokens and learning contextual dependencies through self-supervised objectives. Similar trends have emerged in epigenomics, where models such as EpiAgent [17] extend foundation-model-style pretraining to chromatin accessibility data. More recently, multi-omics foundation models such as SCARF [18] and scTranslator have begun to explore large-scale pretraining for cross-modal representation learning and modality translation. Together, these studies mark an important transition from dataset-specific modeling toward more general-purpose single-cell representations.

Despite this progress, existing single-cell foundation models still face several critical limitations. First, many models are restricted to a single omic modality or a specific modality pair. As a result, they do not provide a unified framework for modeling the full regulatory cascade from chromatin state to transcriptional output and protein-level phenotype. Second, current models often lack a common representation space in which epigenomic, transcriptomic, and proteomic information can be jointly encoded, aligned, and compared. This limits their utility for unpaired integration, missing-modality inference, and holistic cell-state characterization. Third, generative capabilities remain underdeveloped. Some models are primarily optimized for representation learning rather than cross-omics generation, while others adopt autoregressive architectures with causal, unidirectional attention, such as scMulan [19] and TranscriptFormer [20]. Although such designs are effective for sequential data, their unidirectional attention is less naturally aligned with cellular omics profiles, where genes, cis-regulatory elements, and proteins form a jointly dependent set that requires global contextual modeling during generation. Therefore, a unified generative foundation model that can simultaneously support transferable representation learning, multi-omics alignment, and reliable cross-modal generation across epigenomics, transcriptomics, and proteomics remains lacking.

To overcome these limitations, we present **HoloCell**, a unified generative foundation model for holistic cellular modeling across three single-cell omics modalities, namely epigenomics, transcriptomics, and proteomics. HoloCell is anchored on genes and introduces a unified hierarchical tokenization strategy that encodes cis-regulatory elements, genes, and proteins as structured tokens. It employs bidirectional attention and an iterative diffusion-based generation framework. Through staged pretraining on large-scale unimodal and paired multimodal data, HoloCell achieves three core capabilities. First, it provides unified representation by mapping cellular states from multiple omic perspectives into a shared embedding space, thereby capturing cell heterogeneity as an integrated system. Second, it enables unified generation through its iterative diffusion and remasking framework, aligning with a non-autoregressive paradigm to facilitate digital simulation of multi-omic information flow. Third, it supports zero-shot cross-modal inference, allowing diverse downstream tasks such as cross-omics translation without fine-tuning. Extensive experiments demonstrate that HoloCell achieves superior performance across single-omics and multi-omics representation learning, unpaired alignment, and cross-modal generation, providing a flexible and unified foundation model toward the emerging concept of a virtual cell.

## 2 Results

### 2.1 Overview of HoloCell

HoloCell is designed as a unified generative foundation model for holistic cellular modeling across single-cell epigenomics, transcriptomics, and proteomics. Rather than developing separate models for individual modalities or task-specific settings, HoloCell aims to convert heterogeneous omic measurements into a unified sequence representation, allowing a single Transformer encoder to learn unimodal structure, cross-modal correspondence, and generative relationships across omics layers. To achieve this goal, HoloCell is built upon a large-scale single-cell multi-omics corpus and integrates four key technical components: a gene-anchored hierarchical tokenization strategy, a bidirectional Transformer encoder, and an iterative remasking-based generative decoding procedure (Figure 1).

**Figure 1.**
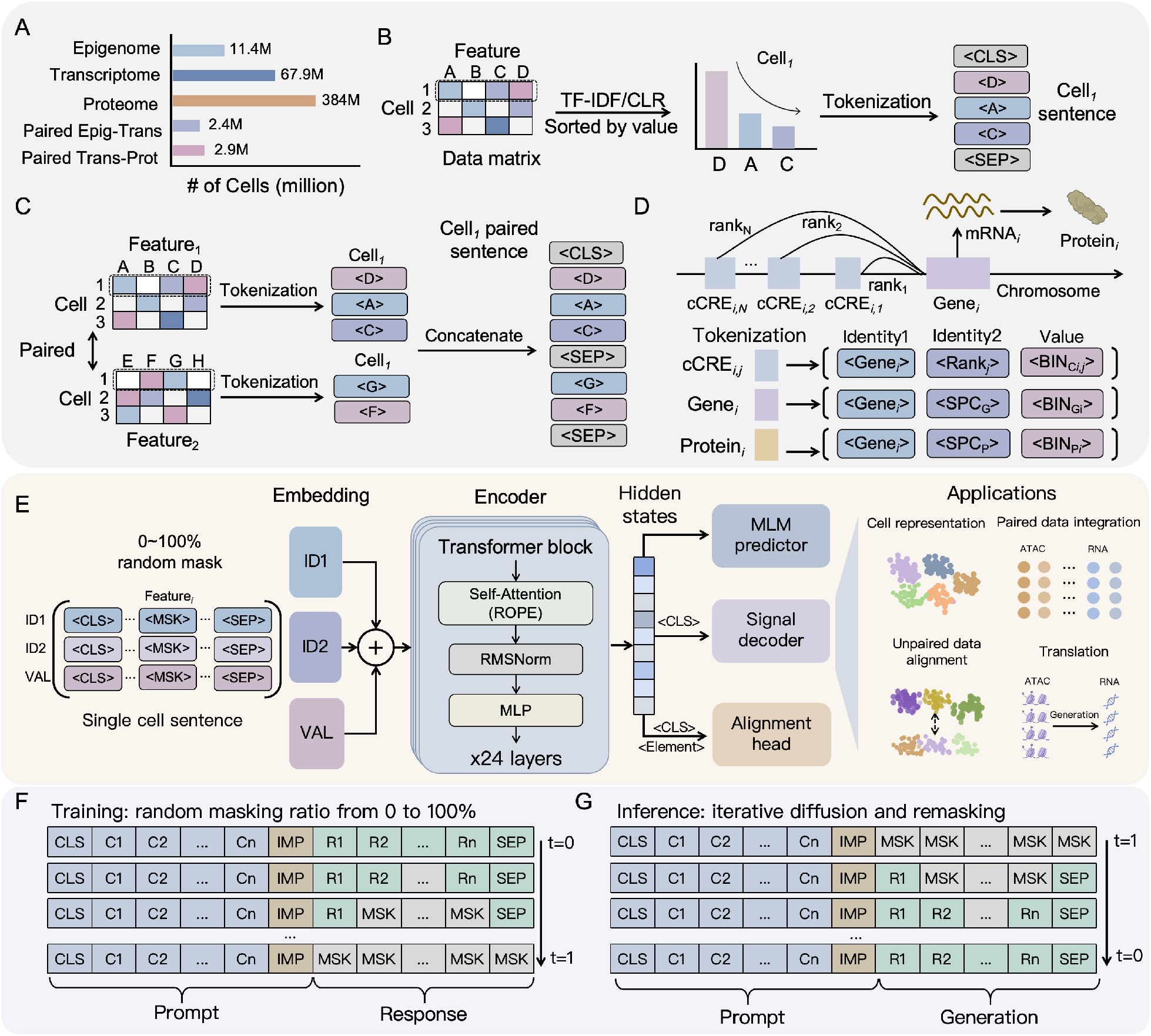
Overview of HoloCell framework. (A) Composition of the HMOC-468M pretraining corpus, comprising 67.9M single-cell transcriptomes, 11.4M single-cell epigenomes, 383.5M single-cell proteomes, 2.4M paired epigenomic-transcriptomic cells, and 2.9M paired transcriptomic-proteomic cells. (B) Unimodal sequence construction pipeline. (C) Paired multi-omics sequence construction via concatenation of aligned unimodal sequences. (D) Unified hierarchical tokenization strategy, mapping each biological element (cCRE, gene, or protein) into a triplet of discrete tokens: identity token, modality or rank token, and value token discretized into 512 bins. ¡SPC¿ denotes the special token representing the modality type. (E) Model architecture of HoloCell, consisting of a triplet embedding layer (identity1, identity2, value), a 24-layer bidirectional Transformer encoder with 20 attention heads and hidden dimension 1,280, and three multi-task output heads: masked prediction heads, signal reconstruction heads, and cell-element alignment heads. (F-G) Iterative diffusion-based generation framework. (F) Post-training stage for generative tasks, featuring dynamic masking ratio *ρ* ~ *U*(0, 1) and contiguous span masking. (G) Inference stage for generative tasks with iterative masked decoding, where the model progressively fills the most confident masked positions until complete generation.

HoloCell is pretrained on the **Human-Multi-Omics-Corpus-468M** (HMOC-468M), which, to our knowledge, is the largest curated human single-cell multi-omics resource assembled to date. This corpus contains approximately 67.9 million single-cell transcriptomes, 11.4 million single-cell epigenomes, and 383.5 million single-cell proteomes, together with over 2.4 million cells with paired epigenomic–transcriptomic measurements and 2.9 million cells with paired transcriptomic–proteomic measurements (Figure 1A). This scale and modality coverage provide the data foundation for learning regulatory and functional relationships across omics layers. For unimodal data, non-zero biological elements are first sorted in descending order according to their processed signal values and then samples them under modality-specific length constraints: 8,192 for epigenomics, 4,096 for transcriptomics, and 1,024 for proteomics. These modality-specific limits are chosen based on corpus-wide length distributions, allowing over 75% of cells to be represented without truncation while preserving major biological signals and controlling computational cost. For paired multi-omics data, unimodal sequences from the same cell are concatenated into a composite sequence, thereby preserving the natural correspondence between modalities within the same biological entity (Figure 1B,C).

To represent heterogeneous omic elements within a shared sequence space, HoloCell implements a gene-anchored hierarchical tokenization strategy (Figure 1D). Specifically, each biological element is represented as a triplet of discrete tokens. The first token specifies the anchored gene identity; the second token indicates either the modality type or, for cCREs, the rank-order distance to its anchor gene; and the third token discretizes the continuous measurement into one of 512 value bins. In this way, feature identity, modality origin, regulatory positional relationship, and quantitative signal intensity are jointly encoded within a unified vocabulary. The design is motivated by biological interpretability and computational efficiency. From a biological perspective, regulatory elements closer to their anchored gene, such as cCREs, are generally associated with stronger regulatory potential, and the hierarchical tokenization naturally captures this positional–functional relationship. From a computational perspective, assigning a single independent token to every gene, cCRE, and protein would result in an excessively large vocabulary, potentially involving over one million candidate tokens, which makes generative prediction inefficient. Unlike Cisformer [21], which decomposes CRE identifiers into digit-level components, HoloCell adopts a biologically anchored hierarchy that couples gene identity with modality and signal values, enabling multi-omic integration into a unified foundation model for both representation learning and diffusion-based generation.

The model backbone is a 24-layer bidirectional Transformer encoder with 20 attention heads and a hidden dimension of 1,280 (Figure 1E). The three token types, namely identity1, identity2, and value, are independently mapped into 1,280-dimensional embeddings, summed, and normalized with RMSNorm to form the final input representation [22]. Unlike autoregressive models that rely on a fixed left-to-right generation order, bidirectional attention allows HoloCell to model dependencies among biological elements using the full cellular context, an approach better aligned with the non-autoregressive modeling paradigm required for holistic single-cell omic interpretation. During pretraining, HoloCell jointly optimizes three complementary objectives. The dynamic masked prediction objective applies span masking with lengths from 1 to 16 and masking ratios uniformly sampled from [0,1], requiring the model to recover masked token triplets from their surrounding context [23]. The signal reconstruction objective encourages the class token (CLS) representation to preserve sufficient global information for reconstructing modality-specific biological signal profiles. The cell–element alignment objective further establishes fine-grained semantic associations between global cell states and individual omic features through contrastive learning [24, 25]. These objectives are combined with equal weighting, enabling the model to capture local feature dependencies, global cellular states, and cell-to-element correspondence within a unified training framework.

Built upon this unified representation and pretraining framework, HoloCell further supports cross-omics generation through an iterative diffusion-inspired remasking strategy (Figure 1F,G). During training, the model receives paired multi-omics sequences in which a subset of target-modality tokens is randomly masked, and it learns to predict the masked target tokens conditioned on the observed source modality and the remaining target context. During inference, generation starts from a fully masked target-modality sequence conditioned on a fully observed source modality. At each iteration, the model predicts all masked positions in parallel and fills a subset of high-confidence tokens according to their prediction probabilities; this procedure is repeated until the target sequence is fully decoded. By progressively refining a masked target sequence through repeated prediction and confidence-based token filling, this process follows the spirit of recent masked diffusion language models, such as LLaDA [26], and resembles the gradual denoising procedure in discrete diffusion models [27, 28]. Importantly, because tokens are predicted in parallel and selected by confidence rather than generated sequentially from left to right, HoloCell avoids imposing an artificial generation order on genes, cis-regulatory elements, or proteins. As a result, HoloCell can perform cross-modal translation, such as epigenome-to-transcriptome generation and transcriptome-to-proteome prediction, without introducing modality-specific architectural modifications.

### 2.2 Representation Learning for Single Omics

A core capability of a single-cell foundation model is to encode individual cells into representations that are informative for downstream biological analysis and transferable across datasets. As illustrated in Figure 2A, existing approaches for single-cell representation learning are often designed for a specific omic modality, such as scGPT for transcriptomics, EpiAgent for epigenomics, and cytoVI for proteomics. These models have contributed substantially to modality-specific analysis, but they typically require separate architectures or training procedures for different data types. By contrast, HoloCell provides a single, unified framework that jointly processes transcriptomic, epigenomic, and proteomic measurements, allowing us to demonstrate for the first time that a single foundation model can learn transferable, biologically informative representations across all three major single-cell omics modalities simultaneously.

**Figure 2.**
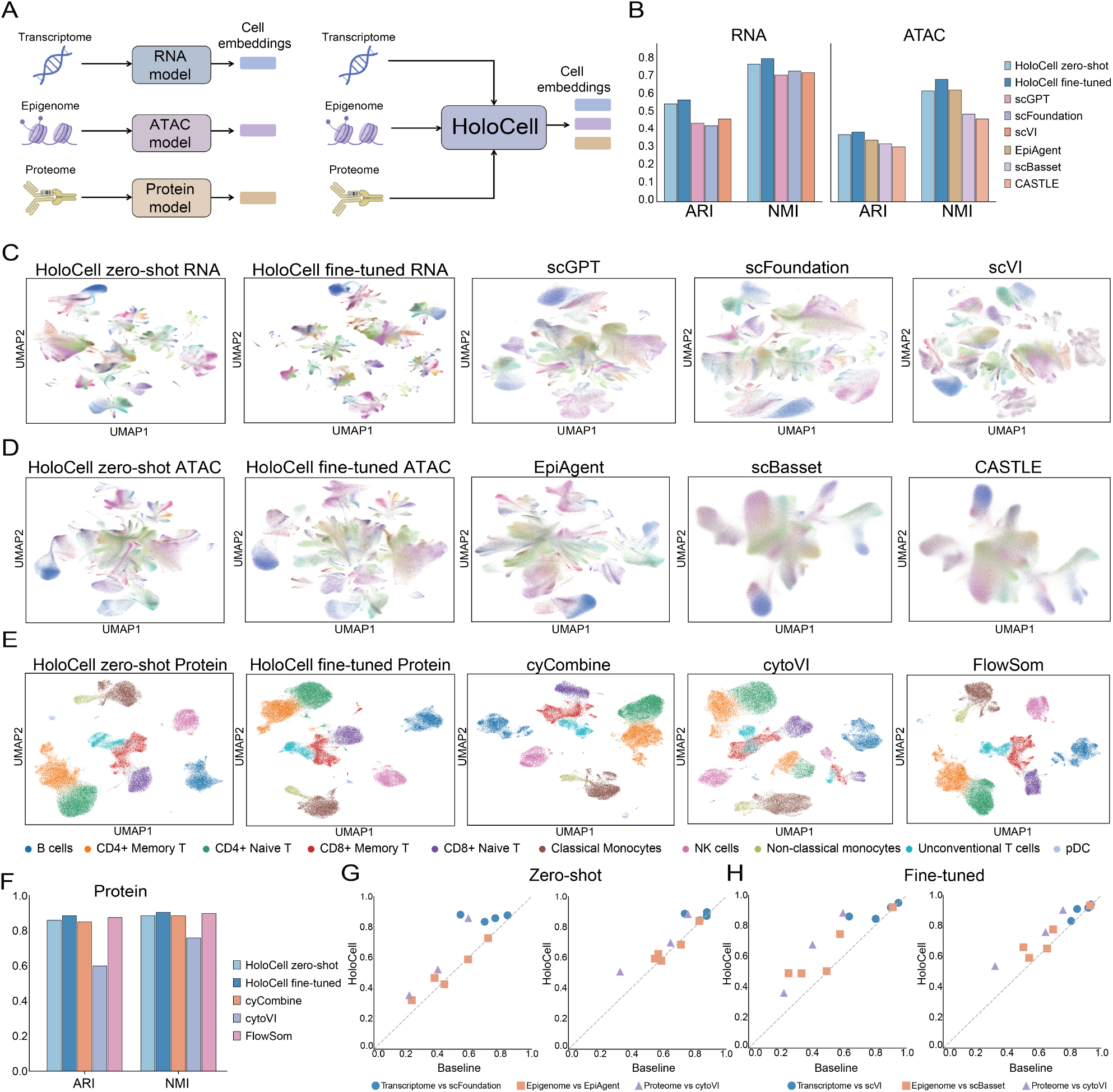
Single-omics representation learning with HoloCell. (A) Schematic illustration comparing traditional modality-specific approaches (separate models for each omics type) with HoloCell’s unified framework that extracts cellular representations across transcriptomics, epigenomics, and proteomics using a single model. (B) Bar charts comparing clustering performance (ARI and NMI) of HoloCell (zero-shot and fine-tuned) against baseline methods on HDMA RNA (left) and ATAC (right) datasets. (C-D) UMAP visualizations of HDMA RNA and ATAC representations colored by cell type annotations. (E) UMAP visualizations of PBMC proteomics data comparing different methods. (F) Bar charts comparing ARI and NMI performance on proteomic Kotliarov2020. (G-H) Scatter plots comparing HoloCell against baseline methods across multiple independent test sets under zero-shot (G) and fine-tuned (H) settings, demonstrating consistent superior performance. Each point represents a dataset-modality pair; points above the diagonal indicate HoloCell outperforming the baseline.

To evaluate this capability, we benchmarked HoloCell under both zero-shot and fine-tuned settings against representative modality-specific methods on independent test datasets. For transcriptomic and epigenomic modalities, we used the Human Development Multiomic Atlas (HDMA) [29], a large-scale single-cell atlas containing chromatin accessibility and gene expression profiles from 817,740 fetal cells across 12 organs. For transcriptomic representation learning, HoloCell was compared with scGPT and scFoundation in the zero-shot setting, as well as scVI, which was fitted on the HDMA dataset. For epigenomic representation learning, we compared HoloCell with EpiAgent in the zero-shot setting, together with scBasset and CASTLE, both of which were trained or fitted on HDMA. Clustering performance was evaluated using Adjusted Rand Index (ARI) [30] and Normalized Mutual Information (NMI) [31] against the ground-truth cell type annotations, which include 134 distinct cell types in HDMA.

HoloCell demonstrated superior performance on both transcriptomic and epigenomic representation learning tasks, demonstrating that the same pretrained model can extract informative cellular embeddings from distinct omic layers (Figure 2B–D). In the transcriptomic setting, zero-shot HoloCell already outperformed the strongest baseline by 18.2% in ARI and 5.3% in NMI. After fine-tuning, the improvement increased to 22.9% in ARI and 9.5% in NMI, indicating that the pretrained representation can be further adapted to dataset-specific cell-state structures. In the epigenomic setting, zero-shot HoloCell achieved an 8.7% ARI improvement over the strongest baseline while maintaining a comparable NMI, and fine-tuning further increased the gains to 13.0% in ARI and 9.5% in NMI. The corresponding UMAP embeddings showed coherent cell-type organization in both RNA and ATAC data, with clearer population separation in RNA, consistent with the higher sparsity and regulatory complexity of chromatin accessibility profiles (Figure 2C,D). Detailed metric values are provided in the supplementary table (Supplementary Table 1).

We next evaluated whether HoloCell can generalize to lower-dimensional proteomic profiles using the large-scale Kotliarov2020 CyTOF dataset [32]. Unlike RNA and ATAC data, proteomic measurements typically contain only tens to hundreds of markers, presenting a representation learning problem with substantially different dimensionality and sparsity. HoloCell was compared with proteomics- and cytometry-oriented methods including cyCombine, cytoVI, and FlowSOM, all of which require fitting on the test dataset. Remarkably, even in a zero-shot setting without any dataset-specific training, HoloCell achieved performance comparable to the strongest baselines. After fine-tuning, HoloCell further outperformed these methods (Figure 2E,F). Together, these results demonstrate that HoloCell provides a broadly applicable representation framework across transcriptomic, epigenomic, and proteomic data, rather than relying on separate modality-specific models for each single-cell assay.

To further examine generalization across datasets, we extended the evaluation to multiple independent test sets, including 4 transcriptomic datasets, 5 epigenomic datasets, and 3 proteomic datasets. In the zero-shot setting, HoloCell was compared with scFoundation for transcriptomics, EpiAgent for epigenomics, and cyCombine for proteomics. In the fine-tuned setting, comparisons were made against scVI for transcriptomics, scBasset for epigenomics, and cyCombine for proteomics. Figures 2G and 2H present scatter plots in which each point corresponds to a dataset-modality pair, with the baseline score on the x-axis and the HoloCell score on the y-axis. Most points fall above the diagonal, indicating that HoloCell often matches or improves upon the corresponding baseline across the evaluated datasets. This trend is observed in both zero-shot and fine-tuned settings, supporting the transferability of the learned representations across different data sources and modalities. Overall, these results suggest that HoloCell provides a flexible and broadly applicable representation learning framework for single-cell transcriptomic, epigenomic, and proteomic data.

### 2.3 Integration of Paired Multi-Omics Data

Beyond learning representations from individual omics layers, an important objective of HoloCell is to integrate paired multi-omics measurements from the same cell and extract a more complete representation of cellular state. In paired single-cell multi-omics data, each modality captures a distinct biological aspect of the same cellular entity: chromatin accessibility reflects regulatory potential, transcript abundance reflects gene activity, and protein abundance provides a closer readout of cellular phenotype. Therefore, an effective paired multiomics model should learn a joint representation that preserves cell-type-resolved biological structure while leveraging complementary information across modalities. We evaluated this capability in paired ATAC–RNA integration, paired RNA–protein integration, and tri-omics ATAC–RNA–protein integration.

We first assessed HoloCell on paired ATAC–RNA datasets (Figure 3A–C). In HSC Ma2020 [33], zero-shot ATAC-only embeddings captured major biological variation but showed limited resolution for closely related populations, whereas RNA-only embeddings yielded clearer cell-type structure. Simultaneous paired input produced a stronger representation, improving ARI/NMI from 0.746/0.767 (RNA alone) and 0.460/0.524 (ATAC alone) to 0.855/0.830 in the zero-shot paired setting—already close to target-fitted baselines. Fine-tuning further improved performance to 0.875/0.847. Similar patterns were observed on PBMC 10xMultiome [34], where HoloCell’s paired zero-shot (0.752/0.811 ARI/NMI) outperformed the strongest dataset-fitted baseline (MOJITOO [35]) by 24.6% in ARI and 5.8% in NMI, with fine-tuning achieving further gains (0.763/0.843). Detailed comparisons on both datasets are provided in Supplementary Tables 2. Together, these results demonstrate that HoloCell integrates paired regulatory and transcriptional signals without dataset-specific optimization, while fine-tuning enables further adaptation to individual datasets.

**Figure 3.**
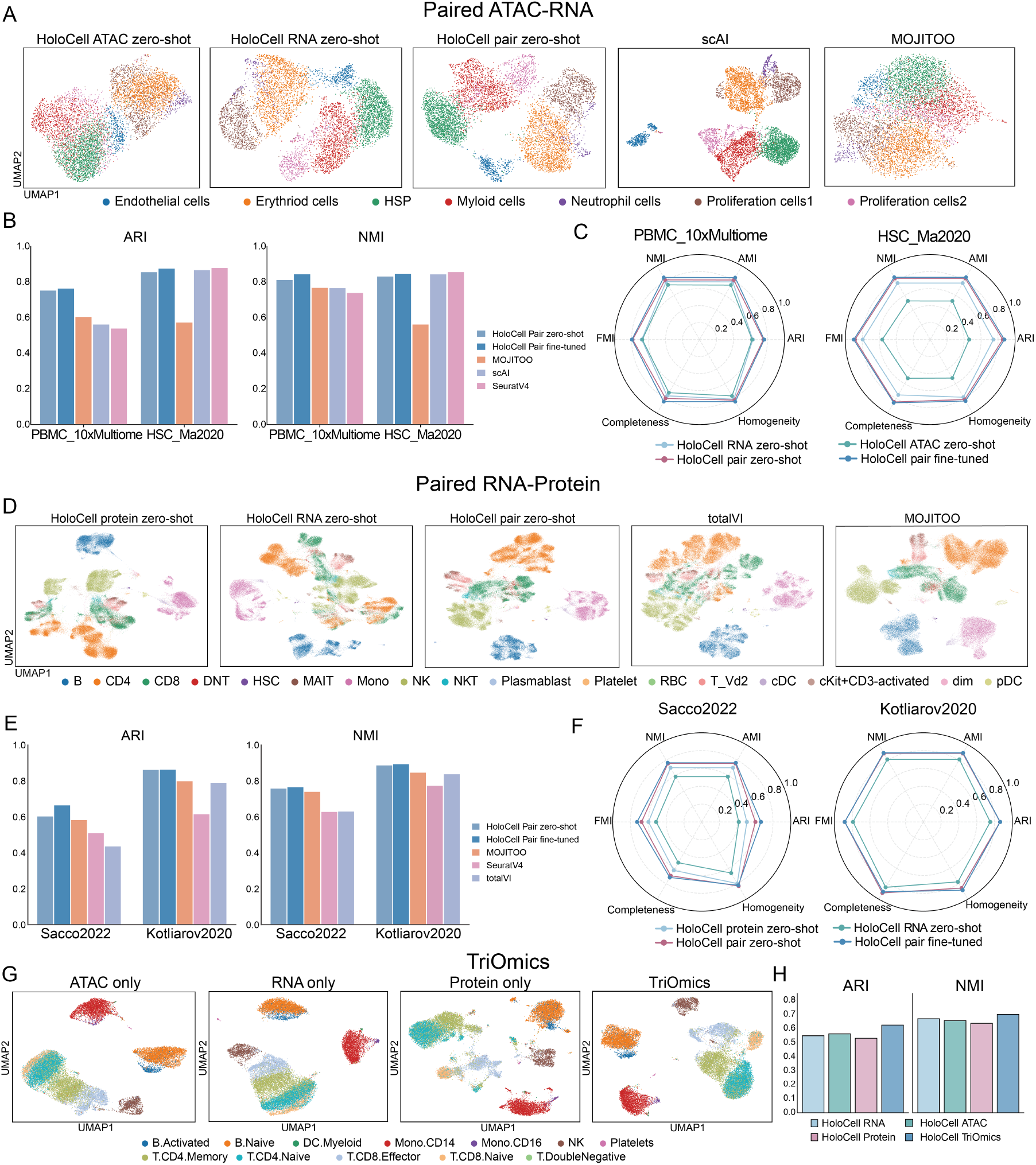
Integration of paired multi-omics data with HoloCell. (A) UMAP visualization of paired ATAC–RNA data (HSC Ma2020) across methods. (B) Clustering performance (ARI/NMI) on PBMC 10xMultiome and HSC Ma2020 for HoloCell (zero-shot/fine-tuned) vs. baseline methods. (C) Radar plots comparing HoloCell representations from ATAC-only, RNA-only, paired zero-shot, and paired fine-tuned inputs. (D) UMAP visualization of paired RNA–protein data (Kotliarov2020) across methods. (E) Clustering performance on Sacco2022 and Kotliarov2020 for RNA–protein integration. (F) Radar plots comparing HoloCell protein-only, RNA-only, paired zero-shot, and paired fine-tuned representations. (G) UMAP visualization of TEA-seq tri-omics data using single-modality and tri-omics inputs. (H) Clustering performance comparison for HoloCell using RNA-only, ATAC-only, protein-only, and tri-omics inputs.

We next evaluated paired RNA–protein integration (Figure 3D–F). On Kotliarov2020 [32], which exhibits pronounced batch effects, HoloCell produced a high-quality paired embedding in the zero-shot setting, preserving major immune populations and yielding a more organized representation than RNA alone. The paired zero-shot embedding achieved ARI/NMI of 0.862/0.887, substantially higher than RNA-only input (0.751/0.809) and comparable to the highly discriminative protein-only representation (0.863/0.889). It also surpassed the strongest baseline, MOJITOO, by 7.9% in ARI and 4.8% in NMI. On Sacco2022 [36], HoloCell paired zero-shot attained 0.602/0.758 ARI/NMI, exceeding MOJITOO (0.582/0.740) by 3.4% in ARI and 2.4% in NMI; fine-tuning further improved these to 0.665/0.766, widening the ARI gain to 14.2%. Paired RNA–protein input provided a clear benefit over RNA alone on both datasets and over protein alone on Sacco2022, demonstrating that HoloCell effectively integrates complementary information from gene expression and surface protein abundance (see Supplementary Table 3 for full results).

Finally, we investigated whether HoloCell can generalize to tri-omics integration. Although HoloCell was trained on paired ATAC–RNA and paired RNA–protein data rather than fully paired ATAC–RNA–protein tri-omics data, it can naturally accept all three modalities through the same unified tokenization and sequence construction framework. We evaluated this zero-shot tri-omics capability on a TEA-seq dataset [37] (Figure 3G,H). ATAC-only and RNA-only representations captured broad immune cell structure but showed limited separation between closely related CD4+ T and CD8+ T populations, while protein-only representations better distinguished these T cell subsets but produced a less organized global manifold. In contrast, the tri-omics representation integrated complementary information from all three biological layers and achieved the best clustering performance, with ARI/NMI of 0.625/0.702. This corresponds to an 11.1% improvement over the best unimodal ARI from ATAC-only input and a 4.6% improvement over the best unimodal NMI from RNA-only input. These results demonstrate that HoloCell can flexibly compose information across modalities and generalize beyond the paired combinations observed during pretraining, highlighting its potential as a unified foundation model for paired and higher-order single-cell multi-omics integration.

### 2.4 Alignment of Unpaired Multi-Omics Data

A key challenge in single-cell multi-omics analysis is aligning different omic modalities when cell-level pairing information is unavailable. In such settings, the goal is to construct a shared representation space in which cells with similar biological states are close across modalities, enabling label transfer from well-characterized RNA datasets to ATAC datasets and facilitating comparative analysis across heterogeneous atlases.

We therefore evaluated whether HoloCell can support unpaired ATAC–RNA alignment. First, HoloCell was adapted from the pretrained model using paired ATAC–RNA data through a CLIP-like contrastive learning objective [38] (Figure 4A). In this training stage, paired ATAC and RNA profiles from the same cell were encoded separately, and their cell embeddings were encouraged to be closer in the shared representation space than non-matched cells. After this contrastive adaptation, HoloCell can be applied to unpaired datasets by independently extracting ATAC and RNA embeddings and then aligning them in the same latent space. Importantly, this inference procedure does not require cell-type labels or cell-level correspondence in the target dataset.

**Figure 4.**
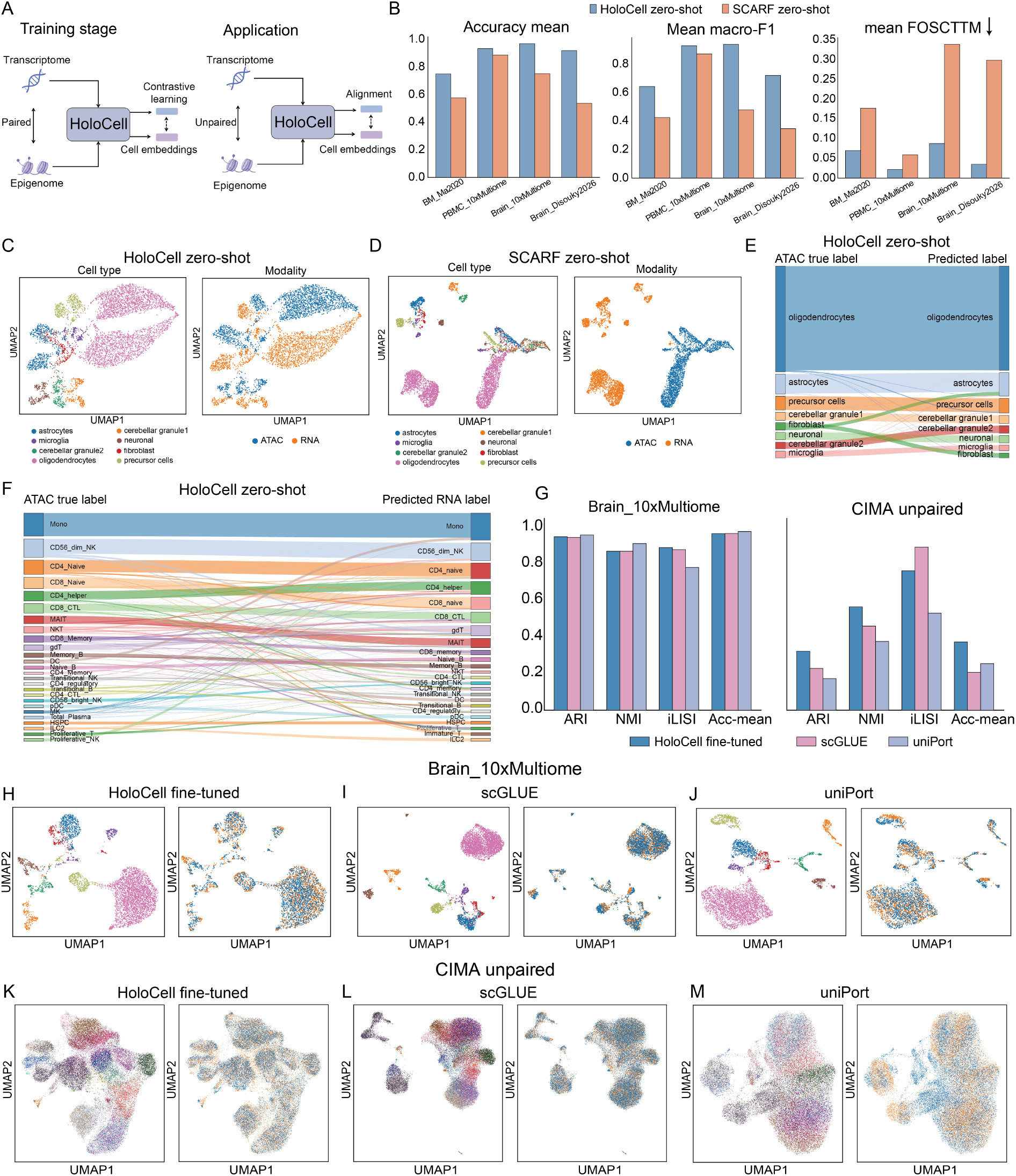
Alignment of unpaired ATAC–RNA data with HoloCell. (A) Schematic of contrastive adaptation on paired ATAC–RNA data and zero-shot application to unpaired ATAC and RNA profiles. (B) Zero-shot comparison between HoloCell and SCARF across paired test datasets using cross-modal label-transfer accuracy, macro-F1, and mean FOSCTTM. (C–D) UMAP visualizations of zero-shot embeddings on Brain 10xMultiome for HoloCell and SCARF, colored by cell type and modality. (E–F) RNA-to-ATAC label-transfer results for HoloCell zero-shot embeddings on Brain 10xMultiome and the CIMA unpaired test set. (G) Quantitative comparison of fine-tuned HoloCell, scGLUE, and uniPort on Brain 10xMultiome and CIMA unpaired data using ARI, NMI, iLISI, and neighbor-based matching accuracy. (H–J) UMAP visualizations on Brain 10xMultiome for HoloCell, scGLUE, and uniPort. (K–M) UMAP visualizations on the CIMA unpaired test set for HoloCell, scGLUE, and uniPort. 12

We first examined the zero-shot alignment ability of HoloCell, using SCARF [18] as a baseline foundation model for ATAC–RNA representation learning. To quantify whether the zero-shot embedding captures cross-modal biological similarity, we evaluated four paired ATAC–RNA test datasets while hiding the pairing information during inference. Three complementary metrics were compared (Figure 4B). The first two metrics are the mean accuracy and mean macro-F1 of bidirectional cross-modal label transfer based on weighted nearest-neighbor voting, where labels from one modality are used to predict labels in the other modality. These metrics evaluate whether cross-modal neighbors share the same biological annotation. The third metric is mean FOSCTTM, which measures the fraction of cells closer than the true paired cell in cross-modal retrieval; lower values indicate better recovery of cell-level matching structure. Across the evaluated datasets, HoloCell achieved higher label-transfer accuracy and macro-F1 than SCARF, while also showing lower FOSCTTM in most datasets, suggesting that its zero-shot embeddings provide a more informative cross-modal semantic space.

A representative visualization on the Brain 10xMultiome dataset [34] is shown in Figures 4C and D. In the zero-shot setting, HoloCell brought ATAC and RNA cells with related cell types closer together while preserving major biological structures. SCARF also captured part of the cell-type organization, but the two modalities remained more separated in the embedding space. To further test the practical utility of the zero-shot alignment, we performed RNA-to-ATAC label transfer using nearest-neighbor voting. On Brain 10xMultiome, HoloCell accurately transferred RNA-derived labels to ATAC cells, achieving an overall accuracy of 97% (Figure 4E). We next applied the same strategy to the more challenging CIMA unpaired test set, which contains 26 immune cell subtypes collected from three donors. Even without target-specific fine-tuning, HoloCell correctly transferred labels for a substantial fraction of ATAC cells, achieving an accuracy of 0.568 (Figure 4F). These results indicate that HoloCell can provide a useful zero-shot initialization for semantic alignment and label transfer in unpaired ATAC–RNA datasets.

To further refine the shared embedding space on target datasets, we introduced an optimal transport (OT)-guided pseudo-pairing strategy followed by contrastive fine-tuning. Specifically, we first computed zero-shot ATAC and RNA embeddings and used optimal transport to identify candidate cross-modal anchors. To reduce the effect of noisy pseudo-pairs, we applied confidence filtering and top-*K* assignment constraints to retain high-confidence anchors. These pseudo-paired cells were then used for contrastive fine-tuning, encouraging cells with similar biological states to move closer across modalities.

We compared fine-tuned HoloCell with scGLUE [12] and uniPort [41] on both a paired Brain 10xMultiome dataset and the unpaired CIMA test set (Figure 4G). We evaluated ARI and NMI to measure cell-type-resolved biological structure, iLISI [42] to quantify local modality mixing, and neighbor-based cross-modal matching accuracy to assess whether cross-modal neighbors share consistent labels. On the relatively simple paired Brain 10xMultiome dataset, all three methods achieved strong performance, with similar ARI, NMI, and neighbor-matching accuracy. On the more complex CIMA unpaired dataset, HoloCell obtained higher ARI, NMI, and neighbor-matching accuracy than the baselines, indicating better preservation of cell-type-discriminative structure. In contrast, scGLUE achieved a slightly higher iLISI, suggesting stronger local modality mixing. This difference indicates that the methods emphasize different aspects of integration: scGLUE promotes more complete modality mixing, whereas HoloCell retains more cell-type-resolved biological structure in this setting.

The UMAP visualizations further support this interpretation. On Brain 10xMultiome, HoloCell, scGLUE, and uniPort all produced relatively well aligned embeddings (Figures 4H– J). On the CIMA unpaired test set, HoloCell better preserved major immune cell populations while maintaining partial ATAC–RNA mixing (Figure 4K). scGLUE showed stronger modality mixing but less distinct separation among some cell-type groups, whereas uniPort exhibited weaker biological organization in this dataset (Figures 4L–M). Together, these results suggest that HoloCell can support unpaired ATAC–RNA alignment by combining zero-shot cross-modal semantic representations with OT-guided contrastive refinement, providing a practical framework for label transfer and biologically structured multi-omics integration.

### 2.5 Zero-shot Generation following the Central Dogma with HoloCell

The central dogma of molecular biology describes the directional flow of biological information from chromatin regulation to RNA transcription and ultimately to protein expression. Following this biological hierarchy, inferring downstream omic layers from upstream measurements is a central task in single-cell multi-omics modeling. In particular, generating transcriptomic profiles from chromatin accessibility and generating proteomic profiles from transcriptomic measurements can help connect regulatory states with cellular phenotypes. HoloCell is, to our knowledge, the first model capable of zero-shot generation along this central-dogma axis across epigenomic, transcriptomic, and proteomic layers. We therefore evaluated its zero-shot cross-modal generation capability in three settings, including ATAC-to-RNA generation, RNA-to-protein generation, and sequential generation following the epigenome–transcriptome–proteome axis.

We first examined ATAC-to-RNA generation on the PBMC 10xMultiome test dataset. Since currently available published methods provide limited support for direct zero-shot transcriptome generation from chromatin accessibility profiles, we used the gene activity score computed by EpiScanpy [43] as a zero-shot baseline. The RNA profiles generated by HoloCell from ATAC inputs were close to the real RNA profiles in the UMAP space (Figure 5A). When colored by data type, the predicted and real transcriptomes showed considerable overlap, suggesting that HoloCell can recover a transcriptome-like manifold from chromatin accessibility alone. When colored by cell type, the generated profiles retained recognizable immune cell populations, indicating that the predicted transcriptomes preserved a degree of cell-type-specific biological structure. In comparison, the gene activity score representation showed a more visible separation from the real RNA profiles (Figure 5B), suggesting that accessibility-derived gene activity scores only partially approximate the measured transcriptomic states.

**Figure 5.**
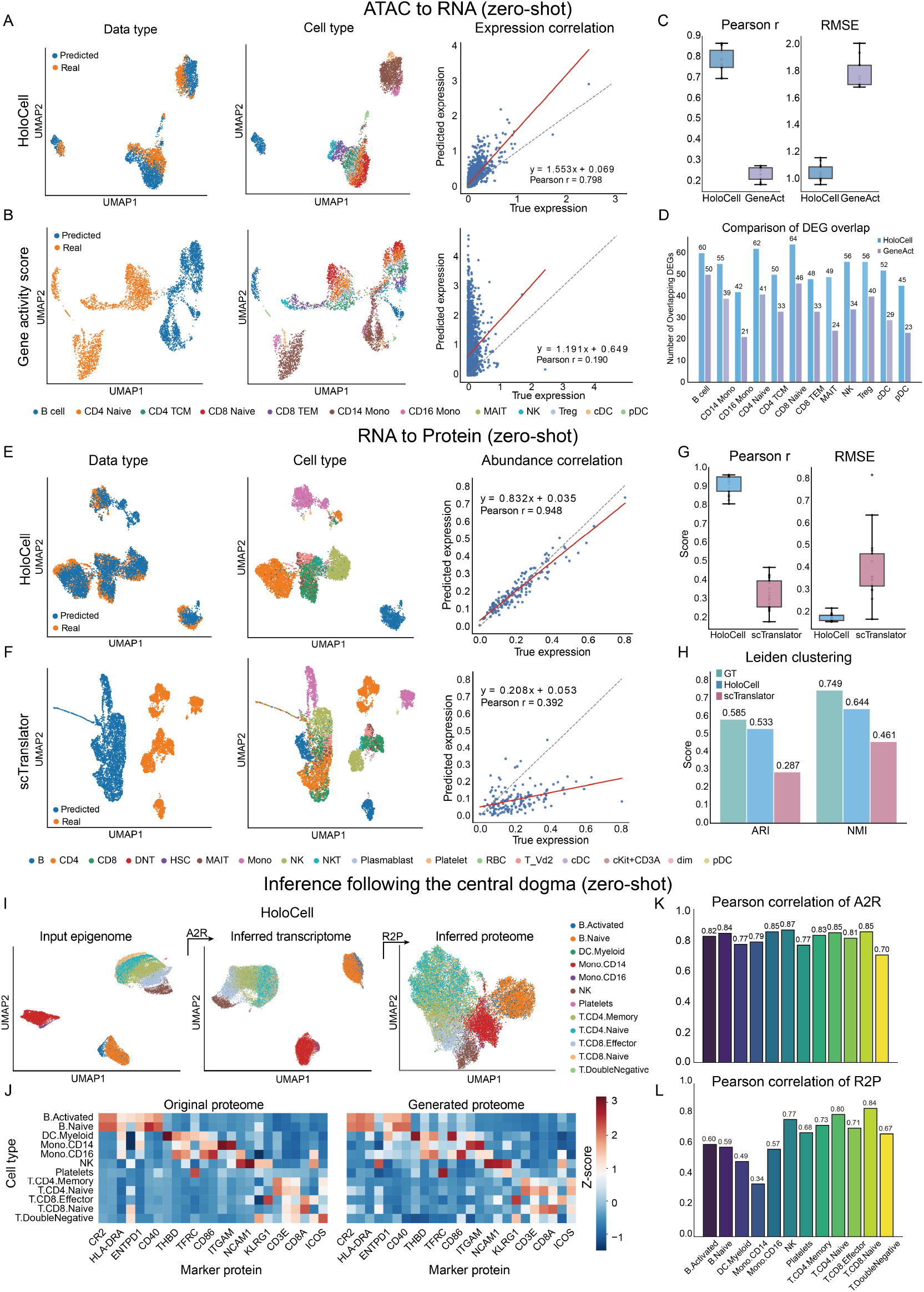
Zero-shot cross-modal generation with HoloCell. (A–D) ATAC-to-RNA generation on PBMC 10xMultiome. Comparison with EpiScanpy gene activity scores. Metrics include UMAP, global correlation, cell-type-wise statistics, and DEG overlap. (E–H) RNA-to-protein generation on Sacco2022. Comparison with scTranslator via UMAP, global and cell-type-wise correlation/RMSE, and clustering metrics. (I–L) Sequential generation on TEA-seq data. Epigenome → transcriptome → proteome. Results include UMAP, marker heatmaps, and cell-type-wise correlations.

We further quantified the ATAC-to-RNA generation results by comparing the predicted and real mean expression values of the top 3,000 highly variable genes across the test dataset. HoloCell achieved a Pearson correlation of 0.798 between predicted and real expression values (Figure 5A), whereas the gene activity score baseline reached a correlation of 0.190 (Figure 5B). To account for cell-type-specific expression patterns, we also computed Pearson correlation and RMSE within each annotated cell type. The cell-type-wise Pearson correlations of HoloCell were mostly distributed between 0.7 and 1.0, while those of the gene activity score baseline were mainly between 0.1 and 0.3 (Figure 5C). The RMSE values of HoloCell were also generally lower, mostly ranging from 0.9 to 1.2, compared with approximately 1.6 to 2.0 for the baseline. We next asked whether the generated transcriptomes could recover cell-type-associated transcriptional programs. The overlap between the top 100 differentially expressed genes identified from generated profiles and those identified from real RNA profiles is shown in Figure 5D. Across the evaluated cell types, HoloCell recovered more real DEGs than the gene activity score baseline, suggesting that the generated transcriptomes better preserve differential expression patterns relevant to cell identity.

We next evaluated RNA-to-protein generation on the Sacco2022 dataset [36]. We used scTranslator [44], a model specifically designed for RNA-to-protein prediction, as the baseline. UMAP visualizations of predicted and real protein profiles generated by HoloCell and scTranslator are compared in Figures 5E and F. For HoloCell, the predicted protein profiles largely followed the manifold of the real protein profiles when colored by data type, while the cell-type-colored visualization showed that major immune populations were retained. For scTranslator, predicted and real protein profiles showed a larger distributional gap, although several cell-type structures were still distinguishable. To make the comparison compatible with the output space of scTranslator, HoloCell predictions were normalized to the range of 0 to 1 before quantitative evaluation. HoloCell achieved a Pearson correlation of 0.948 between predicted and real protein values across all proteins (Figure 5E), whereas scTranslator achieved a correlation of 0.392 (Figure 5F).

Cell-type-wise evaluation showed a similar trend. HoloCell obtained Pearson correlations mostly between 0.8 and 1.0, while scTranslator was mainly distributed between 0.2 and 0.5 (Figure 5G). For RMSE, HoloCell values were concentrated around 0.1 to 0.25, whereas scTranslator showed a broader distribution with higher values in several cell types. We also evaluated whether the generated protein profiles remained useful for downstream clustering. Leiden clustering based on real protein profiles achieved an ARI of 0.585 and an NMI of 0.749 (Figure 5F). HoloCell-generated protein profiles achieved an ARI of 0.533 and an NMI of 0.644, which were lower than the real protein profiles but higher than the scTranslator-generated profiles, whose ARI and NMI were 0.287 and 0.461, respectively. These results suggest that HoloCell-generated protein profiles retain useful cell-state information for down-stream analysis.

Finally, we investigated whether HoloCell can perform sequential generation following the central dogma. Using TEA-seq chromatin accessibility profiles [37] as input, we first generated transcriptomic profiles through ATAC-to-RNA inference and then generated proteomic profiles from the inferred transcriptomes through RNA-to-protein inference. UMAP visualizations of the original epigenomic profiles, inferred transcriptomes, and inferred proteomes, all colored by cell type, are shown in Figure 5I. The inferred transcriptomes preserved the broad cell-type structure, and the generated proteomes also retained major immune cell populations despite being generated from predicted rather than measured RNA profiles. Quantitatively, the first generation stage achieved cell-type-wise Pearson correlations mostly between 0.7 and 0.9 between inferred and real transcriptomes (Figure 5K). In the second generation stage, the correlations varied more across cell types, likely reflecting error propagation from the inferred RNA input (Figure 5L). Nevertheless, the generated protein profiles still captured part of the cell-type-associated proteomic structure. The marker protein heatmaps in Figure 5J further show that several major cell-type-specific marker patterns observed in the real protein data were also present in the generated proteomes. Together, these results indicate that HoloCell can perform zero-shot cross-modal generation across multiple omic layers and can be used to infer downstream biological profiles from upstream measurements.

## 3 Discussion

HoloCell provides a unified foundation model for learning, integrating, and generating single-cell biological profiles across epigenomic, transcriptomic, and proteomic layers. By combining gene-anchored tokenization, large-scale unimodal and paired multimodal pretraining, and bidirectional masked generation, HoloCell supports representation learning, paired and unpaired integration, and cross-modal prediction within a single framework. Across independent test settings, HoloCell produced competitive or improved cell representations in both zero-shot and fine-tuned modes, suggesting that the pretrained model captures transferable cell-state structure rather than merely fitting individual datasets. The paired multi-omics results further show that joint inputs generally improve clustering and manifold organization compared with unimodal inputs, supporting the view that complementary biological layers can be fused into a shared cellular representation.

The generative results further highlight the advantage of this unified design. HoloCell recovered transcriptome-like manifolds from ATAC profiles and protein-like manifolds from RNA profiles without requiring task-specific architectures. These observations suggest that masked diffusion-style decoding can model cross-modal biological dependencies in a way that overcomes the constraints of sequential generation, in contrast to purely autoregressive approaches. The sequential TEA-seq analysis also indicates that HoloCell can compose multiple generation steps along the epigenome–transcriptome–proteome axis.

These findings support three broader implications. First, a common token space anchored on genes can serve as a practical bridge between heterogeneous omic measurements. The same biological coordinate system can represent nearby cis-regulatory elements, gene expression, and protein abundance, improving interpretability while allowing the Transformer backbone to share parameters across modalities. Second, large-scale pretraining on both unimodal and paired data appears useful for generalization. Unimodal data provide scale and broad coverage, whereas paired data teach the model cross-modal correspondences that can later be used for integration, label transfer, and conditional generation. Third, generative objectives expand the role of single-cell foundation models beyond embedding extraction. A model that can infer missing modalities or impute incomplete profiles may help reduce experimental cost, support atlas harmonization, and prioritize hypotheses about regulatory programs and surface phenotypes.

HoloCell also suggests a path toward more complete cellular simulators. Rather than treating RNA, chromatin accessibility, and protein abundance as separate analysis products, a holistic model can represent them as coupled observations of the same latent biological state. This does not remove the need for mechanistic validation, but it provides a scalable statistical substrate for generating hypotheses across biological layers.

Several limitations remain. First, the model is trained primarily on human datasets, so its behavior on non-human species, rare perturbation states, and poorly sampled tissues requires systematic evaluation. Second, although HoloCell can operate on unpaired data, the strongest cross-modal signals still come from paired measurements. Performance may degrade when the target domain contains cell states absent from the paired pretraining corpus. Third, generation quality is currently evaluated mainly through distributional, cell-type-wise, and marker-level metrics. Future studies should include more causal or perturbational validation to determine whether generated modalities preserve regulatory relationships that are biologically actionable. Fourth, the present framework represents cCREs through gene-anchored rank tokens, which is efficient and interpretable but may simplify distal enhancer logic, three-dimensional chromatin interactions, and context-specific regulatory wiring.

Future work should extend HoloCell in several directions. First, expanding the pretraining corpus to additional omic layers and non-human species may improve its applicability beyond the current epigenomic, transcriptomic, and proteomic settings. Second, the generative framework can be further developed for within-modality imputation, enabling recovery of missing or sparsely observed signals within the same omic layer. Third, HoloCell may provide a useful foundation for perturbation response prediction by modeling how cellular states change under genetic, chemical, or environmental perturbations. Finally, the learned cross-modal and cell–element relationships could be explored for gene regulatory network construction and disease-associated regulatory inference, providing a bridge between generative modeling and mechanistic biological interpretation.

## 4 Conclusion

HoloCell establishes a generative foundation model that treats the three major single-cell omic layers, namely epigenomics, transcriptomics, and proteomics, as coupled views of a shared latent cellular state. Through unified hierarchical tokenization anchored on genes, combined with large-scale pretraining on over 468 million single-cell profiles and 425 billion tokens, the model learns transferable representations that preserve cell-type-resolved biological structure across modalities without requiring dataset-specific adaptation. A distinctive strength of HoloCell is its bidirectional iterative diffusion generation framework, which overcomes the constraints of fixed-order causality in modeling genes, cis-regulatory elements, and proteins. This design enables zero-shot cross-modal translation, such as predicting transcriptomes from chromatin accessibility and proteomes from transcriptomes, while avoiding the biologically implausible sequential assumptions of autoregressive models.

By unifying representation learning, paired and unpaired integration, and cross-modal generation within a single architecture, HoloCell moves beyond modality-specific tools toward a holistic cellular simulator. This capability reduces the experimental burden of measuring every layer in every cell, facilitates the construction of multi-modal cell atlases, and opens new avenues for data-driven discovery of cross-omic regulatory programs. As a versatile foundation model, HoloCell represents a concrete step toward the emerging vision of a virtual cell, where diverse biological readouts are generated and interpreted from a common latent representation of cellular identity and state.

## 5 Methods

### 5.1 Data Collection and Preprocessing

To support the large-scale pre-training of HoloCell, we curated a comprehensive multi-omics corpus, designated as the Human-Multi-Omics-Corpus-468M (HMOC-468M), by integrating data from published literature and public repositories, including CELLxGENE [45], CIMA [39], Human-scATAC-Corpus [46], SPDB [47], and scMMO-atlas [48]. HMOC-468M represents the largest and most comprehensive human single-cell multi-omics resource to date, encompassing 67,933,274 single-cell transcriptomes, 11,351,597 single-cell epigenomes, and 383,477,079 single-cell proteomes. Furthermore, HMOC-468M includes paired multi-omics data consisting of 2,443,679 cells with simultaneous epigenomic and transcriptomic profiling, as well as 2,897,667 cells with paired transcriptomic and proteomic measurements.

#### Single-cell transcriptomics

We collected human single-cell transcriptomic profiles from the CELLxGENE database [45] and the CIMA database [39]. Low-quality cells expressing fewer than 1,000 genes or fewer than 3,000 unique molecular identifiers (UMIs) were excluded, yielding an expression matrix of 67,933,274 cells across 19,238 protein-coding genes. To prevent housekeeping genes from masking subtle expression patterns of low-abundance transcription factors [49], we applied a Term Frequency-Inverse Document Frequency (TF-IDF) normalization followed by a log1p transformation with a scaling factor of 10,000.

#### Single-cell epigenomics

We curated single-cell epigenomic data for 11,351,597 cells across 1,355,445 candidate cis-regulatory elements (cCREs) from the Human-scATAC-Corpus database [46] and published literatures [50, 51, 52]. The reference cCREs were derived from a previously established foundation model for single-cell epigenomics [17]. Similar to the transcriptomic data, the raw chromatin accessibility matrix was normalized using the same global TF-IDF transformation strategy.

#### Single-cell proteomics

We compiled single-cell proteomic profiles for 383,477,079 cells from the SPDB database [47], comprising 380,141,177 cells generated via CyTOF [53] and 3,335,902 cells profiled by CITE-seq [54]. For CyTOF data, we applied an arcsinh transformation with a cofactor of 5 to stabilize variance. For CITE-seq antibody-derived tag (ADT) counts, we employed a modified centered log-ratio (CLR) transformation to account for compositionality and sequencing depth variations.

#### Single-cell multi-omics

To support the cross-omics integration and translation, we further curated single-cell paired multi-omics datasets capable of simultaneously profiling multiple modalities within the same individual cell, which primarily fall into two major categories: joint single-cell transcriptomic-epigenomic and transcriptomic-proteomic profiling. Beyond the scMMO-atlas database [48], we expanded our data collection by integrating publicly accessible datasets from published studies, including the Human Development Multiomic Atlas (HDMA) [29] and the human hippocampal neurogenesis atlas [55]. We finally yielded a total of 2,443,679 single-cell transcriptomic-epigenomic cells, alongside 2,897,667 cells characterized by transcriptomic and proteomic measurements. To ensure technical consistency across the entire corpus, the constituent modalities within these paired multi-omics datasets were preprocessed and normalized strictly using their respective single-modality normalization frameworks detailed in the preceding sections.

### 5.2 Tokenization and Sentence Construction

As summarized in Figures 1D, HoloCell simultaneously considers both the identity and the quantitative value of each biological element during tokenization.

#### Hierarchical Tokenization

To address the large vocabulary size induced by the vast number of cCREs (approximately 1.35 × 10^6^), we adopt a gene-anchored hierarchical tokenization strategy. Let Gene_*i*_ denote the *i*-th gene with expression value *G*_*i*_, and Protein_*i*_ denote the protein encoded by Gene_*i*_ with abundance *P*_*i*_. For epigenomic elements, let cCRE_*ij*_ denote the *j*-th closest regulatory element to Gene_*i*_, ranked by genomic distance, with accessibility value *C*_*ij*_.

We define the tokenization function Tokenizer(·) that maps each biological element to a triplet of discrete tokens. For a gene, its expression value *G*_*i*_ is discretized via binning; for a cCRE, its accessibility value *C*_*ij*_ is similarly binned; for a protein, its abundance *P*_*i*_ is binned accordingly. Formally:

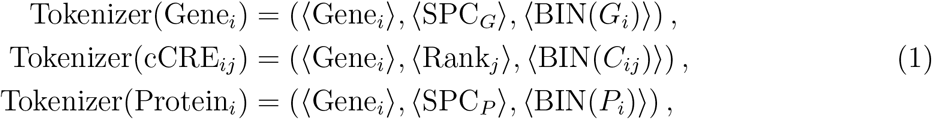

where ⟨SPC_*G*_⟩ and ⟨SPC_*P*_ ⟩ are special tokens distinguishing transcriptomic and proteomic modalities, respectively. BIN(·) denotes a discretization function that maps continuous values (e.g., *G*_*i*_, *C*_*ij*_, *P*_*i*_) into one of *K* = 512 bins.

#### Sentence Construction

For a given cell in a single modality, let ℰ = {*e*_1_, *e*_2_, …, *e*_*n*_} denote the set of non-zero elements sorted in descending order of their processed values. After sampling with a maximum length constraint *L* (where *L* = 8192 for epigenomics, 4096 for transcriptomics, and 1024 for proteomics), we obtain a subset:

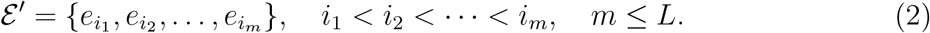

Let the tokenized sequence of the selected elements be

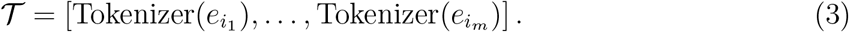

The unimodal cell sentence is then constructed as:

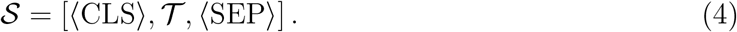

For paired multi-omics data, let *T* ^(*A*)^, *T* ^(*R*)^, and *T* ^(*P*)^ denote the tokenized sequences (without ⟨CLS⟩ or ⟨SEP⟩) for epigenomics, transcriptomics, and proteomics, respectively. The final sentence is constructed by concatenating a single starting ⟨CLS⟩, followed by the token sequence of each modality separated by ⟨SEP⟩, and ending with a final ⟨SEP⟩:

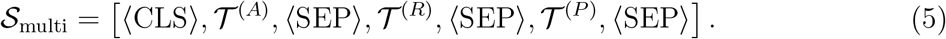

Unless otherwise specified, we adopt a fixed concatenation order following the central dogma of molecular biology: epigenomics (*T*^(*A*)^), followed by transcriptomics (*T*^(*R*)^), and finally proteomics (*T* ^(*P*)^). This ordering reflects the natural flow of genetic information from chromatin accessibility to RNA expression and protein translation. While HoloCell’s bidirectional attention mechanism theoretically supports arbitrary modality permutations, we utilize this biologically informed canonical order during pretraining to align with established regulatory hierarchies.

### 5.3 Model Architecture

Figure 1E presents the overall architecture of HoloCell, which is based on a bidirectional Transformer encoder tailored for multi-omics sequence modeling. The model consists of a triplet embedding layer, a stack of Transformer encoder layers, and multiple task-specific output heads.

#### Embedding Layer

Let **x**^(1)^, **x**^(2)^, **x**^(*v*)^ ∈ ℕ^*T*^ denote the identity1, identity2, and value token sequences of length *T*. These sequences are mapped into continuous embeddings via three independent embedding functions:

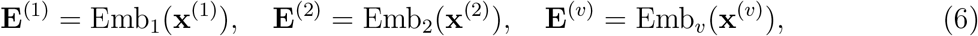

where each embedding lies in ℝ^*T* ×*d*^ with hidden dimension *d* = 1280.

The final input representation is obtained via element-wise summation followed by normalization:

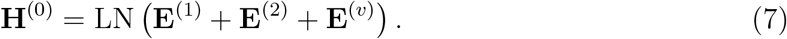

#### Transformer Encoder

The encoder consists of *L* = 24 stacked Transformer layers. Each layer follows a pre-normalization residual structure and is defined as:

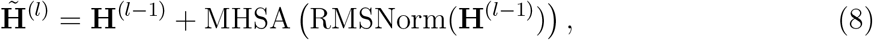

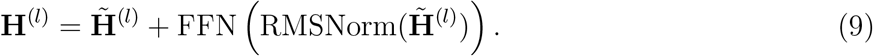

#### Multi-Head Self-Attention

The multi-head self-attention (MHSA) module uses *h* = 20 attention heads. For each head *k*, the attention operation is defined as:

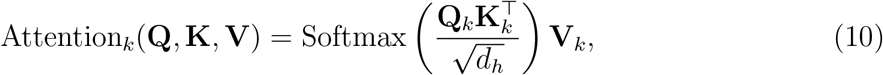

where *d*_*h*_ = *d/h* = 64 is the per-head dimension. The outputs of all heads are concatenated and linearly projected:

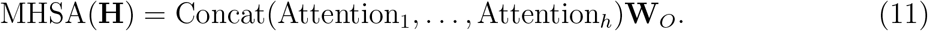

Rotary positional embeddings [56] are applied to the query and key projections to encode positional information, and FlashAttention-2 [57] is used for efficient computation over long sequences.

#### Feed-Forward Network

Each Transformer layer contains a feed-forward network with hidden dimension *d*_ffn_ = 5120. We adopt a gated linear unit (GLU) formulation [58]:

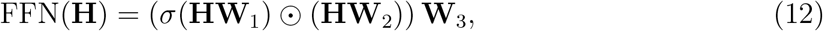

where *σ*(·) denotes the SiLU activation function, ⊙ is element-wise multiplication, and 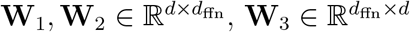.

#### Output Representations

After *L* layers, the final hidden states are:

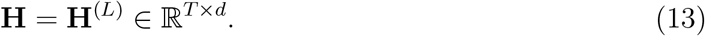

The first token representation **h**_CLS_ = **H**_1_ is used as the global cell representation, while the remaining token representations are used for element-level predictions.

### 5.4 Pretraining Objectives

HoloCell adopts a multi-task learning framework with three complementary objectives: dynamic masked prediction, signal reconstruction, and cell-element alignment.

#### Dynamic Masked Prediction

This task trains the model to predict masked biological elements conditioned on their context. Unlike standard BERT-style masking with a fixed ratio, we employ a dynamic strategy where the masking ratio is uniformly sampled from [0, 1] and masked positions form contiguous spans (lengths 1 to 16). For each masked position *t* ∈ ℳ, the entire triplet is replaced with a special ⟨MASK⟩ token.

Let **H**_*t*_ ∈ ℝ^*D*^ denote the Transformer encoder’s output hidden state at position *t*. Three independent linear prediction heads map **H**_*t*_ to logits over the respective vocabularies:

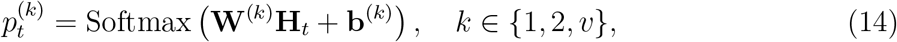

where 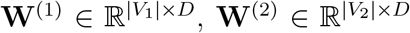, and 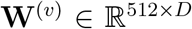. The masked prediction loss is the sum of cross-entropy losses over all masked positions and all three dimensions:

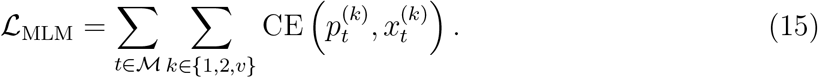

#### Signal Reconstruction

This task learns global cell representations by reconstructing the entire signal profile from the ⟨CLS⟩ token. Let **h**_CLS_ = **H**_1_ ∈ ℝ^*D*^. For each modality *m* ∈ {*A, R, P*} (*A*: epigenomics, *R*: transcriptomics, *P*: proteomics), an MLP-based decoder *f* ^(*m*)^ maps **h**_CLS_ to a reconstructed signal vector:

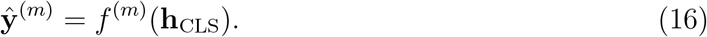

The reconstruction loss combines binary cross-entropy for epigenomics and mean squared error for transcriptomics and proteomics:

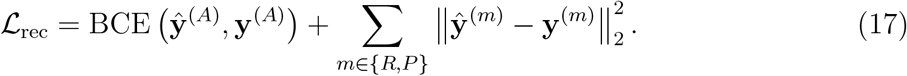

#### Cell-Element Alignment

This task establishes semantic associations between cells and individual biological elements via contrastive learning. For an element *i*, let **h**_*i*_ ∈ ℝ^*D*^ be its representation. The alignment score *s*_*i*_ is computed by an MLP *g*:

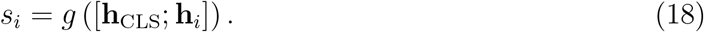

For epigenomics, *s*_*i*_ ∈ [0, 1] predicts the probability that cCRE *i* is accessible, and the loss is binary cross-entropy. For transcriptomics and proteomics, *s*_*i*_ predicts the continuous expression value, and the loss is mean squared error:

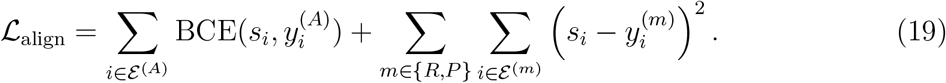

#### Overall Objective

The three losses are jointly optimized via weighted summation:

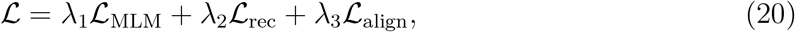

where we set *λ*_1_ = 1.0, *λ*_2_ = 1.0, and *λ*_3_ = 1.0.

### 5.5 Post-training Strategies

#### Cross-modal generation training

Let *s* and *t* denote a source and a target modality, with *s, t* ∈ {*A, R, P*}. Given a paired cell with source sequence *S*^(*s*)^ and target sequence *S*^(*t*)^, we construct a conditional sequence:

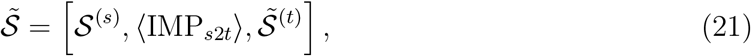

where ⟨IMP_*s*2*t*_⟩ is a task-specific transition token (IMP standing for imputation) and 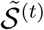 is obtained by dynamically masking the target sequence. Only masked target positions contribute to the conditional generation loss:

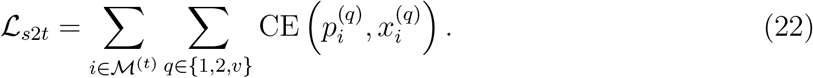

In this work, we used ATAC-to-RNA (*s* = *A, t* = *R*) and RNA-to-protein (*s* = *R, t* = *P*) translation.

#### Diffusion-style iterative generation inference

At inference time, HoloCell performs non-autoregressive masked decoding. Given an observed source sequence *S*^(*s*)^, the target sequence is initialized as fully masked:

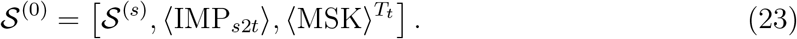

At iteration *r*, the model outputs token distributions for each masked position *i* ∈ ℳ_*r*_:

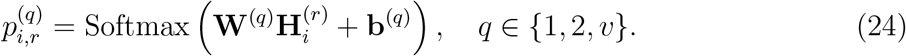

The predicted triplet and its confidence are:

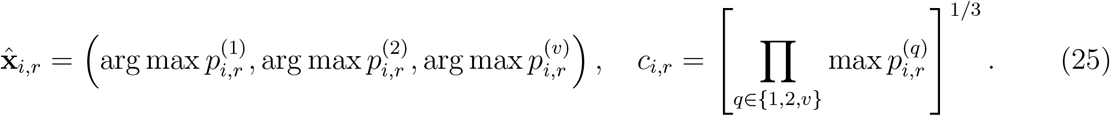

A subset of the most confident masked positions is selected according to a predefined un-masking schedule:

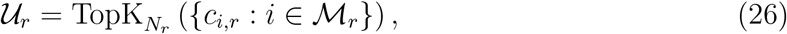

and the sequence is updated as:

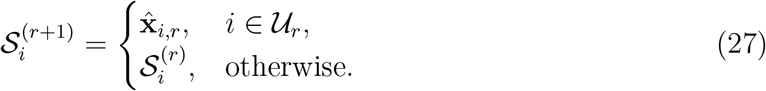

This procedure repeats until ℳ_*r*_ = ∅. Because positions are decoded by confidence rather than rigid sequential order, the inference process aligns with the non-autoregressive paradigm required for single-cell molecular profiling.

#### Multi-omics alignment with contrastive adaptation

We adopt a CLIP-like contrastive learning framework [38]. Given a batch of *N* paired cells, we obtain modality-specific cell representations:

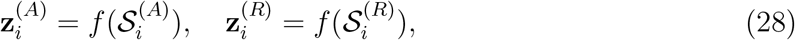

which are *ℓ*_2_-normalized. The cross-modal similarity with temperature *τ* is:

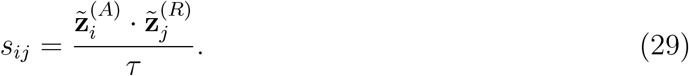

The symmetric InfoNCE objective is:

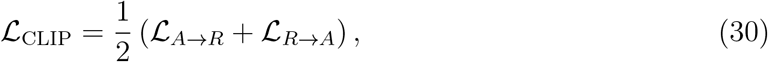

where 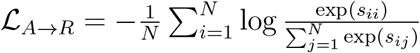, and ℒ_*R*→*A*_ is defined symmetrically.

### 5.6 Fine-tuning Strategies

#### Representation fine-tuning

For cell-representation benchmarks, HoloCell is fine-tuned using only the signal reconstruction objective ℒ_rec_. For single-omics RNA and ATAC, the learning rate is 5 × 10^−5^ with batch size 64 for 3 epochs. For proteomics, the same learning rate is used with batch size 1024. For paired multi-omics, the tokenized paired sequence is used with learning rate 5 × 10^−5^ and batch size 64 for 3 epochs. After fine-tuning, the ⟨CLS⟩ hidden state is extracted as the cell embedding.

#### Contrastive fine-tuning for paired alignment

For paired ATAC–RNA or RNA–protein alignment tasks, HoloCell is fine-tuned with ℒ_CLIP_. Paired cells are encoded independently by modality, with true pairs as positives and other cells in the mini-batch as negatives. Only the top two Transformer layers are updated.

### 5.7 Unpaired alignment with OT-guided pseudo-pairs

For unpaired ATAC–RNA alignment, we first extract zero-shot cell embeddings. Let **Z**^(*A*)^ and **Z**^(*R*)^ denote the *ℓ*_2_-normalized embeddings. The transport cost is defined as cosine distance:

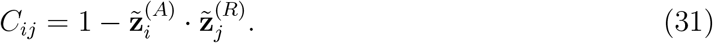

An optimal transport coupling is estimated by solving the entropy-regularized problem [40]:

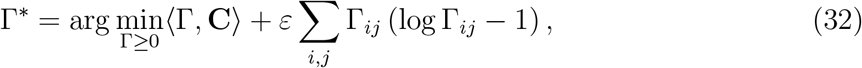

subject to uniform marginal constraints. The coupling is row-normalized to obtain assignment probabilities *P*_*ij*_. An ATAC cell *i* is retained only if max_*j*_ *P*_*ij*_ ≥ 10*/n*_*R*_. For each retained cell, the top-ranked unused RNA cell is selected as its pseudo-pair. The resulting pseudo-pairs are tokenized and used for contrastive fine-tuning with the symmetric InfoNCE loss for 10 epochs with equivalent batch size 64.

### 5.8 Benchmarking and statistical analysis

#### Cell representation, clustering, and visualization

For HoloCell, cell representations were obtained from the final ⟨CLS⟩ hidden state. For baseline methods, we used the latent representation or embedding recommended by each method. Clustering was performed with the Leiden algorithm [59] implemented in `scanpy`. To make clustering scores comparable across methods, the Leiden resolution parameter was selected by binary search so that the number of predicted clusters matched the number of ground-truth cell-type labels. If an exact match was not reachable, we used the resolution producing the closest cluster number. UMAP visualizations were generated using the default `scanpy` parameters [60].

Clustering accuracy was evaluated using Adjusted Rand Index (ARI) and Normalized Mutual Information (NMI), computed with `scikit-learn`. Given a predicted partition *U* and a ground-truth partition *V*, NMI is defined as

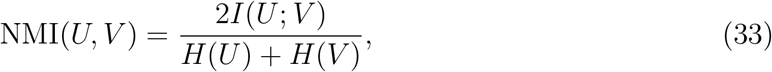

where *I*(*U*; *V*) is mutual information and *H*(·) is entropy. For ARI, let *n*_*ij*_ be the c ontingencytable count between predicted cluster *i* and true label *j, a*_*i*_ = ∑_*j*_ *n*_*ij*_, *b*_*j*_ =∑_*i*_ *n*_*ij*_, and *n* = ∑_*i,j*_ *n*_*ij*_. ARI is computed as [30]

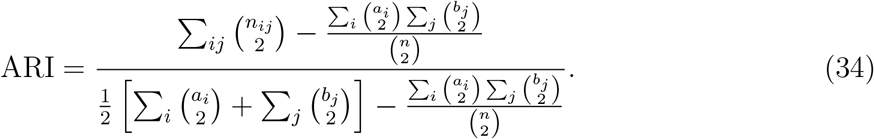

#### Multi-omics alignment metrics

For ATAC–RNA alignment evaluation, ATAC and RNA embeddings were extracted separately and *ℓ*_2_-normalized. Cross-modal label transfer was performed by weighted nearest-neighbor voting with *K* = 10 neighbors. For a query cell *i* from modality *s* and reference cells from modality *t*, the neighbor set *N*_10_(*i*) was selected by cosine similarity. The predicted label was computed as

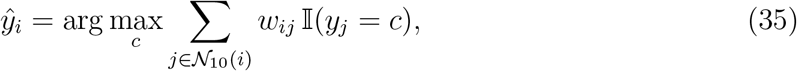

where the weights are obtained by a softmax over cosine similarities:

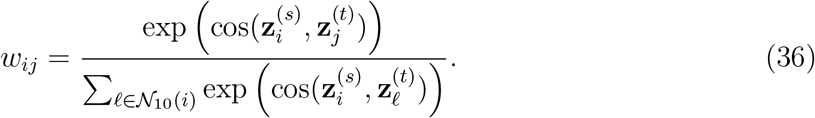

Label transfer was performed in both directions, ATAC-to-RNA and RNA-to-ATAC. The reported accuracy and macro-F1 were averaged across the two directions:

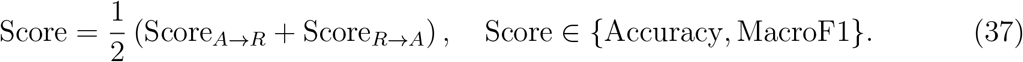

Modality mixing was quantified using iLISI computed by the `scib` implementation, with modality identity used as the batch variable. Higher iLISI values indicate stronger local mixing between modalities, whereas lower values indicate stronger modality separation.

For paired ATAC–RNA test datasets, we also computed FOSCTTM (Fraction of Samples Closer Than the True Match). For query cell *i* and its true paired reference cell *i*, let

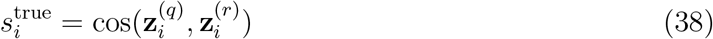

be the true-pair similarity. The number of reference cells closer than the true match is

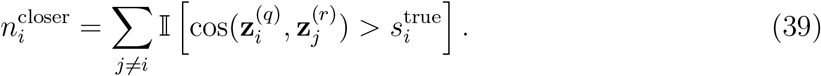

The per-cell and mean FOSCTTM scores are

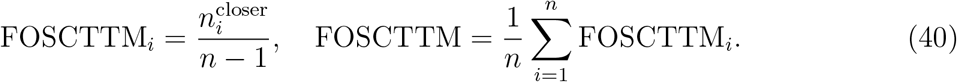

A lower FOSCTTM indicates better recovery of cell-level cross-modal matching, with 0 corresponding to perfect retrieval of the true paired cell.

#### Cross-modal generation metrics

Generated profiles were evaluated against matched ground-truth profiles on a common feature set. For a generated vector **ŷ** and its real target vector **y** with *M* evaluated features, Pearson correlation was computed as

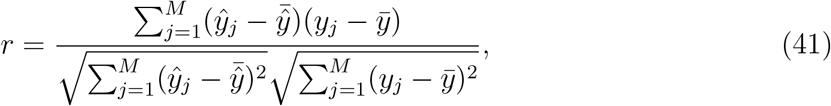

and root mean squared error (RMSE) was computed as

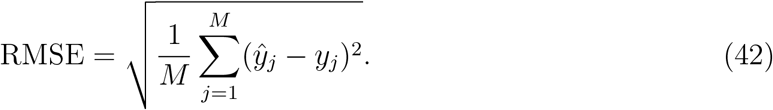

These metrics were reported either globally across the evaluated feature set or within each annotated cell type. For cell-type-wise evaluation, the same formulas were applied after restricting cells to a given annotation group. When differential-expression recovery was evaluated, differentially expressed genes or proteins were ranked separately from generated and real profiles for each cell type, and the overlap between the top-ranked marker sets was reported.

## 6 Data and Code Availability

The pretraining and fine-tuning datasets used in this study are being consolidated and will be made publicly available upon final release. The HoloCell model weights, source code for pretraining, fine-tuning, and inference, as well as comprehensive documentation, are currently under preparation and will be open-sourced at https://github.com/bjzgcai/HoloCell following manuscript revision.

---

**Table 1.**
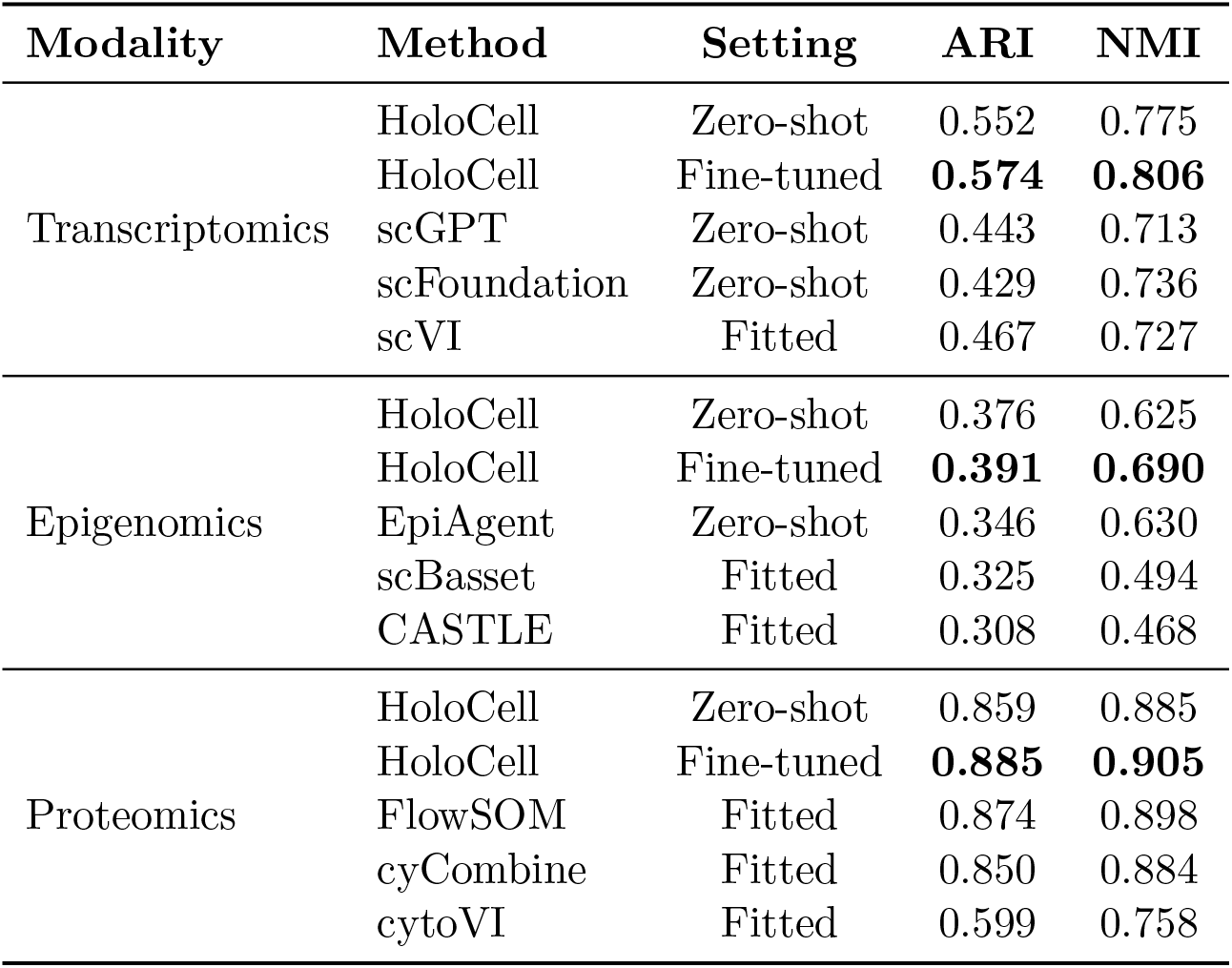
Supplementary Table S1. Performance comparison of HoloCell and baseline methods on single-omics representation learning tasks. ARI and NMI were used to evaluate the agreement between clustering results and cell-type annotations. HoloCell was evaluated in both zero-shot and fine-tuned settings, whereas baseline methods were fitted or applied according to their standard workflows.

**Table 2.** Supplementary Table S2. Performance comparison of HoloCell and baseline methods on paired ATAC-RNA integration tasks (PBMC 10xmultiome and HSC Ma2020). ARI and NMI were used to evaluate the agreement between clustering results and cell-type annotations. Baseline methods were fitted according to their standard workflows.

| Dataset | Method | Setting | ARI | NMI |
| --- | --- | --- | --- | --- |
| PBMC_10xmultiome | HoloCell | ATAC zero-shot | 0.620 | 0.743 |
|  | HoloCell | RNA zero-shot | 0.629 | 0.786 |
|  | HoloCell | Pair zero-shot | 0.752 | 0.811 |
|  | HoloCell | Pair fine-tuned | **0.763** | **0.843** |
|  | MOJITOO | Fitted | 0.604 | 0.767 |
|  | SeuratV4 | Fitted | 0.539 | 0.738 |
|  | scAI | Fitted | 0.561 | 0.766 |
| HSC_Ma2020 | HoloCell | ATAC zero-shot | 0.460 | 0.524 |
|  | HoloCell | RNA zero-shot | 0.746 | 0.767 |
|  | HoloCell | Pair zero-shot | 0.855 | 0.830 |
|  | HoloCell | Pair fine-tuned | 0.875 | 0.847 |
|  | MOJITOO | Fitted | 0.573 | 0.561 |
|  | SeuratV4 | Fitted | **0.878** | **0.856** |
|  | scAI | Fitted | 0.866 | 0.843 |

**Table 3.** Supplementary Table S3. Performance comparison of HoloCell and baseline methods on paired RNA-Protein integration tasks (Sacco2022 and Kotliarov2020). ARI and NMI were used to evaluate the agreement between clustering results and cell-type annotations. Baseline methods were fitted according to their standard workflows.

| Dataset | Method | Setting | ARI | NMI |
| --- | --- | --- | --- | --- |
| Sacco2022 | HoloCell | RNA zero-shot | 0.417 | 0.587 |
|  | HoloCell | Protein zero-shot | 0.507 | 0.702 |
|  | HoloCell | Pair zero-shot | 0.602 | 0.758 |
|  | HoloCell | Pair fine-tuned | **0.665** | **0.766** |
|  | MOJITOO | Fitted | 0.582 | 0.740 |
|  | SeuratV4 | Fitted | 0.510 | 0.628 |
|  | totalVI | Fitted | 0.435 | 0.630 |
| Kotliarov2020 | HoloCell | RNA zero-shot | 0.751 | 0.809 |
|  | HoloCell | Protein zero-shot | 0.863 | 0.889 |
|  | HoloCell | Pair zero-shot | 0.862 | 0.887 |
|  | HoloCell | Pair fine-tuned | **0.863** | **0.894** |
|  | MOJITOO | Fitted | 0.799 | 0.846 |
|  | SeuratV4 | Fitted | 0.614 | 0.774 |
|  | totalVI | Fitted | 0.791 | 0.837 |

**Table 4.** Supplementary Table S4. Zero-shot performance of HoloCell on TEA-seq tri-omics integration (ATAC, RNA, and Protein). ARI and NMI were used to evaluate the agreement between clustering results and cell-type annotations.

| Method | Setting | ARI | NMI |
| --- | --- | --- | --- |
| HoloCell | ATAC zero-shot | 0.563 | 0.658 |
| HoloCell | RNA zero-shot | 0.550 | 0.671 |
| HoloCell | Protein zero-shot | 0.532 | 0.638 |
| HoloCell | Tri-omics zero-shot | **0.625** | **0.702** |

